# Developing forebrain synapses are uniquely vulnerable to sleep loss

**DOI:** 10.1101/2023.11.06.565853

**Authors:** Sean M. Gay, Elissavet Chartampila, Julia S. Lord, Sawyer Grizzard, Tekla Maisashvili, Michael Ye, Natalie K. Barker, Angie L. Mordant, C. Allie Mills, Laura E. Herring, Graham H. Diering

## Abstract

Sleep is an essential behavior that supports lifelong brain health and cognition. Neuronal synapses are a major target for restorative sleep function and a locus of dysfunction in response to sleep deprivation (SD). Synapse density is highly dynamic during development, becoming stabilized with maturation to adulthood, suggesting sleep exerts distinct synaptic functions between development and adulthood. Importantly, problems with sleep are common in neurodevelopmental disorders including autism spectrum disorder (ASD). Moreover, early life sleep disruption in animal models causes long lasting changes in adult behavior. Different plasticity engaged during sleep necessarily implies that developing and adult synapses will show differential vulnerability to SD. To investigate distinct sleep functions and mechanisms of vulnerability to SD across development, we systematically examined the behavioral and molecular responses to acute SD between juvenile (P21-28), adolescent (P42-49) and adult (P70-100) mice of both sexes. Compared to adults, juveniles lack robust adaptations to SD, precipitating cognitive deficits in the novel object recognition test. Subcellular fractionation, combined with proteome and phosphoproteome analysis revealed the developing synapse is profoundly vulnerable to SD, whereas adults exhibit comparative resilience. SD in juveniles, and not older mice, aberrantly drives induction of synapse potentiation, synaptogenesis, and expression of peri-neuronal nets. Our analysis further reveals the developing synapse as a convergent node between vulnerability to SD and ASD genetic risk. Together, our systematic analysis supports a distinct developmental function of sleep and reveals how sleep disruption impacts key aspects of brain development, providing mechanistic insights for ASD susceptibility.

**Significance Statement:** Sleep is a fundamental pillar of lifelong health. Sleep disruption is commonly associated with neurodevelopmental conditions including autism spectrum disorder (ASD) and schizophrenia. Therefore, understanding the vulnerabilities associated with developmental sleep loss is an essential research question. Here we systemically examine the molecular and behavioral consequence of sleep deprivation (SD) in developing and adult mice. Compared to adults, developing mice show absent or blunted adaptive responses to SD, and heightened sensitivity to SD-induced cognitive deficits. Our molecular analysis indicates sleep plays an important role in key aspects of brain development including synaptogenesis, and that the effects of SD converge on nodes of genetic risk for ASD. This study provides new insights into the role of sleep in healthy brain development.

## Introduction

Sleep is a lifelong, essential, and conserved process that supports health and cognition^1^. Sleep behavior and function are controlled by circadian and homeostatic mechanisms that promote sleep at the appropriate time of day and in response to time spent awake^2^. However, what is actually being restored by sleep remains mysterious. While behavioral switching between wake/sleep states is controlled by discrete neuronal populations^3^, the need for sleep is broadly expressed across the brain^4,5^. The neuronal synapse has emerged as one major conserved target for the restorative benefits of sleep and a distributed locus for the accumulation of sleep need^6^. In a previous study, we combined sub-cellular fractionation and quantitative proteomic and phosphoproteomic analyses to demonstrate that synapses of the adult mammalian forebrain undergo profound remodeling throughout the wake/sleep cycle, involving hundreds of proteins and thousands of phosphorylations^7^. This work built upon literature suggesting that part of the restorative basis of sleep involved a broad, but selective, weakening of excitatory synapses through a synaptic plasticity type known as homeostatic scaling down^6,8^. Subsequent multi-omics approaches demonstrated that while the synaptic transcriptome (localized mRNA) is regulated largely as a function of the circadian rhythm, the synaptic proteome and phosphoproteome are regulated almost exclusively as a function of wake/sleep states^9,10^, highlighting synapses as a cellular interface between circadian and sleep-based mechanisms. In a contemporary study, Wang and colleagues demonstrated that brain wide “sleep need” was associated with net accumulation of protein phosphorylation, most notably on synaptic proteins, providing a novel distributed synaptic biochemical substrate for sleep need^5^. This work is in line with the discovery and elucidation of sleep promoting kinases such as SIK3 and CaMKIIβ, and development of a “phosphorylation hypothesis” of sleep^11–15^.

Sleep physiology and synapse density are highly dynamic during postnatal development^16,17^, leading to a critical unanswered question: does sleep perform the same beneficial function throughout the lifespan? Sleep disruption is commonly associated with neurodevelopmental conditions such as Autism Spectrum Disorder (ASD) and Schizophrenia^18–21^. Therefore, understanding the unique vulnerabilities associated with sleep loss during development becomes an essential question of biomedical research. Importantly, sleep disruption during sensitive periods of brain development is causal in long-lasting changes in adult behavior in animal models^22–25^. While a broad homeostatic scaling-down of synapses has been demonstrated in the mature nervous system^7,8^, evidence for this plasticity type was not found in the immature cortex^26^. During development, sleep supports the formation, maturation, and pruning of neuronal synapses, and consolidation of critical period ocular dominance plasticity^27–32^. Together, these points lead to the clear hypothesis that sleep performs unique functions supporting brain and synapse maturation, and further imply that sleep disruption will drive distinct consequences across life stages. This idea is supported by recent mathematical modeling concluding that sleep, across species, plays a completely distinct role during early development supporting brain growth, transitioning to homeostatic functions with maturation^33^.

To test this idea, and to identify nodes of vulnerability to sleep loss in the developing brain, we systematically analyzed the molecular and behavioral effects of acute 4hr or 6hr SD (SD4/6), starting from lights on (ZT0) in wild-type juvenile (postnatal day P21-28), adolescent (P42-49), and adult (P70-100) mice of both sexes. Building from the success of recent multi-omics approaches^5,9,10^, we characterize the effects of SD on the forebrain synapse proteome and phosphoproteome. We corroborate prior findings that daily dynamics of the adult synapse proteome and phosphoproteome are almost exclusively driven by wake/sleep states^9,10^, but further demonstrate that the developing synapse is uniquely and profoundly affected by SD. SD in juveniles, and not adults, drives immediate impacts on key aspects of brain growth such as synaptogenesis, and drives biochemical changes that overlap with nodes of genetic risk for ASD. Our findings provide strong support for an essential role of sleep in healthy brain development.

## Results

During sleep, adult rats and mice exhibit broad dephosphorylation of synaptic AMPA receptor subunit GluA1 at S845 and S831^7,34^, established markers of homeostatic scaling-down and long-term depression^35,36^. The density of forebrain synapses undergoes dramatic changes during postnatal maturation: synaptogenesis, pruning, and stabilization^17^, motivating the idea that sleep may mediate distinct synaptic processes across periods of synapse maturation and stabilization. To test this idea, we systematically examined wake/sleep levels of phospho-GluA1 S845/S831 in mouse forebrain post-synaptic density (PSD) fractions across ages P21-P70. Samples were collected 4hrs into the wake or sleep phase, ZT16 (W4) or ZT4 (S4) respectively, as done previously^7^. Excitingly, unlike prior observations in adults, W4/S4 phospho-GluA1 was completely unaltered from P21-P35, whereas “adult-like” GluA1 dephosphorylation clearly emerged at P42, and was consistent into adulthood at P70 (Fig. 1A and B). This straightforward dataset reveals P42 as an important transitional age, and provides support for the existence of distinct synaptic processes engaged during sleep between development and adulthood^33^. Informed from these results, and motivated by evidence that sleep plays an important role in brain development^23–25^, we hypothesized that developing and adult mice would exhibit distinct consequences and unique vulnerabilities to sleep deprivation (SD). To critically test this hypothesis, in the remainder of this study we engaged in a systematic examination of the behavioral and molecular consequences of acute SD in three developmental ages: Juveniles (P21-P28), Adolescents (P42-P49), and Adults (P70-100) (Fig. 1C).

**Figure 1.**
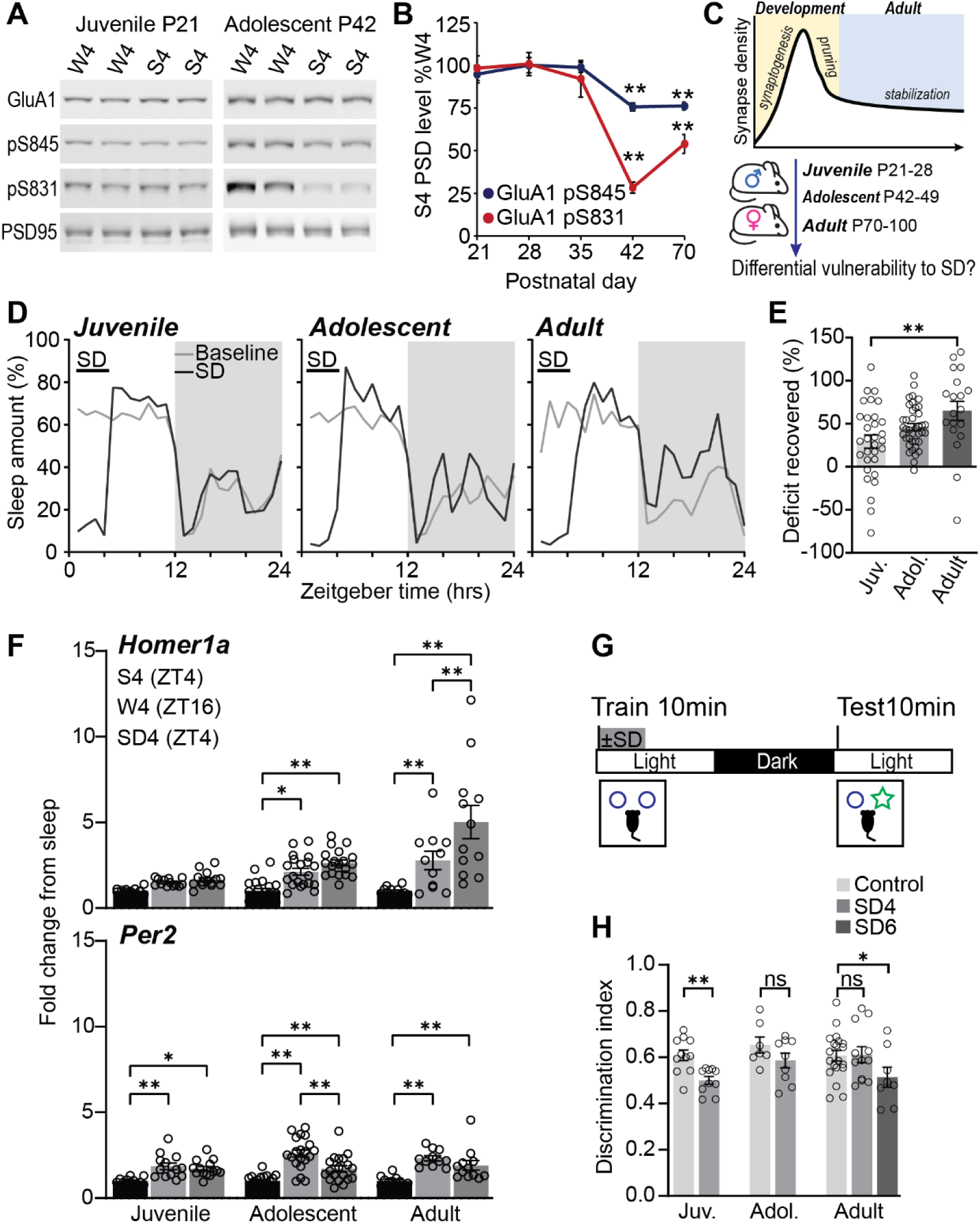
Divergent responses to SD in juveniles, adolescents, and adults. (**A and B**) Western blot analysis of phosphorylated GluA1 S845/S831 in forebrain PSD fractions isolated during wake (W4, ZT16) or sleep (S4, ZT4) from mice aged P21-P70. Phospho-S845/S831 normalized to total GluA1. Mice P42 and older show a significant dephosphorylation during sleep: S4 compared to W4: **P<0.01 1-way ANOVA, Tukey’s correction. (**C**) Schematic: developmental synapse dynamics predicts differential responses to SD. (**D**) 24hr sleep trace during uninterrupted baseline or during 4hr SD treatment (SD4: ZT0-4) and subsequent recovery (ZT4-24) from juvenile (P28), adolescent (P42), or adult mice (P100). Shaded box indicates dark phase. (**E**) % Sleep deficit recovered between ages. Deficit calculated based on sleep lost compared to baseline during SD4 treatment ZT0-4; recovery calculated from increased sleep over baseline from ZT4-24. % Sleep deficit recovered is significantly greater in adults than juveniles. **P<0.01, 1-way ANOVA, Tukey’s correction. (**F**) *Homer1a* and *Per2* transcripts measured using qPCR in whole forebrain from juvenile (P21), adolescent (P42), or adult mice (P100). Samples collected during the sleep phase (ZT4), wake phase (ZT16), or immediately following SD4 (ZT4). *P<0.05, **P<0.01, 1-way ANOVA, Tukey’s correction. (**G and H**) Juvenile (P28), adolescent (P42), or adult mice (P100) subjected to the novel object recognition task ±SD. Training occurs ZT0-2 followed by uninterrupted control or 4-6hr SD treatment, testing occurs 24hrs after training. Novel object memory performance is completely impaired by SD4 in juveniles, memory impaired by SD6 in adults. *P<0.05, **P<0.01, ns: not significant, Student’s t-test. Data includes both sexes. Error bars, mean ±SEM.

### Developing mice exhibit heightened vulnerability and lack robust adaptations to sleep deprivation compared to adults

Adult mammals respond to SD through physiological, behavioral, and molecular adaptations. An increase in non-rapid eye movement sleep slow-wave activity (NREM-SWA) measured by electroencephalography (EEG) over daily baseline has become the most established physiological marker of sleep need^2^. Nelson and colleagues used EEG to systemically examine baseline and recovery sleep following acute SD in mice aged P15-P87. They report that the established SD response of increased NREM-SWA over baseline is absent in juvenile mice and becomes apparent as mice mature to P42 and beyond^37^, suggesting the physiological response to acute SD undergoes important and underappreciated aspects of maturation during these periods of development. At the behavioral and molecular levels SD induces sleep rebound behavior, an increase in subsequent sleep amount during recovery, and broad induction of immediate early gene and homeostatic scaling factor *Homer1a*^38–40^, referred to as “the molecular marker of sleep need”. Surprisingly, whether these well characterized adaptations to SD in adults are differentially expressed during development is not well described.

We used a piezoelectric based non-invasive home-cage sleep recording system to systematically measure uninterrupted baseline sleep and rebound responses following acute 4hr SD (SD4), starting from lights on (ZT0-4), in juvenile, adolescent and adult mice. Daily baseline sleep was highly comparable at each age, with the expected nocturnal pattern of greater sleep amounts during the light phase (Fig. 1D). Although SD4 resulted in comparable levels of total sleep deficit (Fig. S1A, B), distinct SD4 recovery responses were seen at each age. Compared to adults, the developmental ages showed a greater gain of sleep during the remaining 8hrs of the light phase, described as light-phase rebound (Fig. S1C). In contrast, adults showed the expected^2^ significant increase in dark phase sleep following SD4 referred to as dark-phase rebound. Strikingly, dark-phase rebound was completely absent in juveniles and severely blunted in adolescents (Fig. 1D, and Fig. S1A, D). Collectively, despite the small amount of light-phase rebound exhibited by the developmental ages, adult mice with their robust dark-phase rebound were able to recover the majority of sleep loss within 24hrs, whereas developing mice retained the majority of accumulated sleep deficit (Fig. 1E). Next, we measured forebrain transcripts of homeostatic scaling factor *Homer1a* and circadian clock component *Per2* in undisturbed mice 4hrs into the sleep phase (S4/ZT4), wake phase (W4/ZT16), or immediately following SD4 (ZT4). In response to SD4, adults showed the expected^38^ robust induction of *Homer1a* compared to S4, a response that was also completely absent in juvenile mice, and severely blunted in adolescents. In contrast, comparable SD-induction of *Per2*^38^ was observed in all three ages (Fig. 1F), suggesting that circadian adaptations to sleep loss are intact by the juvenile age. Together with the previous systematic EEG analysis of the effects of SD across development^37^, our findings suggest the absence of well characterized behavioral, physiological, and molecular adaptations to sleep loss may expose juvenile mice to greater negative consequences of SD, negatively affecting cognition and brain maturation. Accordingly, memory performance in the novel object recognition (NOR) task was completely impaired in juvenile mice treated by SD4 immediately following training in comparison to undisturbed controls (Fig. 1G, H, and Fig. S1F). In contrast, memory performance in adolescents and adults was resilient to SD4; extended SD6 impaired memory performance in adults, showing that NOR performance is sleep sensitive across ages (Fig. 1H and Fig. S1F). These findings clearly demonstrate that juvenile mice display heightened vulnerability to the consequences of SD compared to older mice.

### The synapse proteome of juvenile, adolescent, and adult mice are distinctly affected by SD

To identify synaptic mechanisms underlying developmental vulnerability to SD, we measured the immediate effects of SD on the developing and adult forebrain synapse proteome (Fig. 2) and phosphoproteome (Fig. 3), using subcellular fractionation and tandem-mass-tag (TMT) quantitative mass spectrometry. Mice were maintained on a 12:12hr light/dark cycle. Juveniles, adolescents, and adults were sacrificed undisturbed during the sleep phase (S4; ZT4), wake phase (W4; ZT16), or immediately following SD4 (ZT4); additional adults were sacrificed immediately following SD6 (ZT6) with matching undisturbed sleep (S6/ZT6) and wake conditions (W6/ZT18). We then isolated the forebrain synapse fraction, commonly referred to as the “post-synaptic density” (PSD), for analysis (Fig. 2A and B)^7^. We identified and quantified ∼5,000 proteins in forebrain synapse fractions from each age. Importantly, the majority of synapse proteins detected (4,485) were identified in all three ages (Fig. 2C), suggesting that we are able to reasonably compare the synaptic response to SD across the ages tested here.

**Figure 2.**
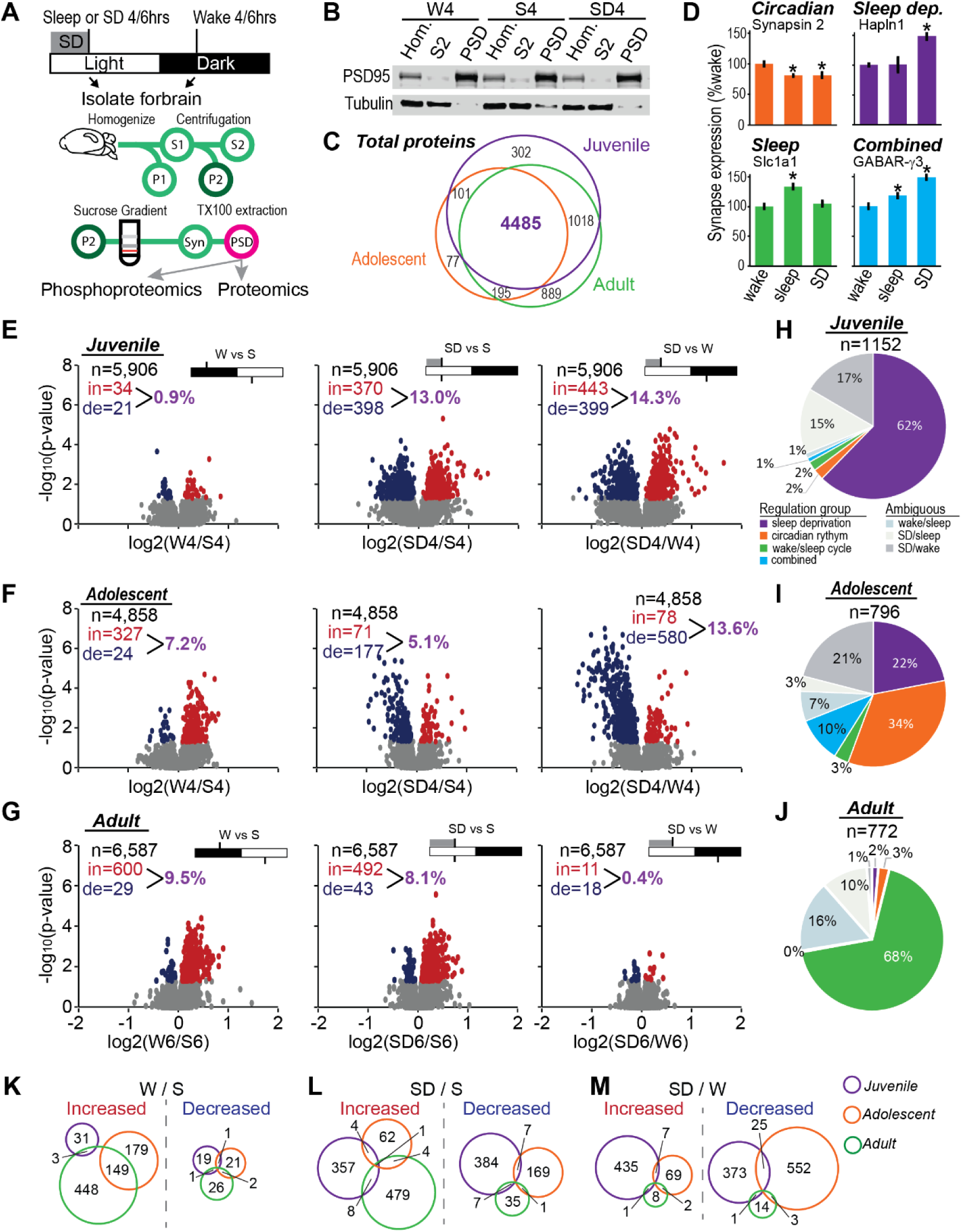
The juvenile synapse proteome is highly vulnerable to SD, transitions to wake/sleep regulation in adults. (**A**) Experimental design. Isolated forebrain synapse fractions (PSD) from juvenile (P21), adolescent (P42), or adult mice (P100) analyzed by quantitative mass spectrometry. 4 mice pooled per sample; samples prepared in quadruplicate. Juveniles and adolescents: W4/S4/SD4; adults: W6/S6/SD6 conditions (**B**) Representative western blot from juvenile cohorts demonstrating enrichment of synapse scaffold protein PSD95 in the PSD fraction from wake, sleep, or SD conditions. (**C**) Venn diagram depicts high overlap in the proteins detected in the synapse fractions from juveniles, adolescents or adults. (**D**) Examples of juvenile synapse proteins regulated by circadian rhythm (circadian), wake/sleep cycle (sleep), selective response to SD (sleep dep.) or combined. *P<0.05, Student’s *t* test with Bonferroni correction. (**E-G**) Volcano plots showing changes in synapse proteins in wake/sleep, SD/sleep, and SD/wake comparisons from juvenile, adolescent, or adult mice. N: number of proteins quantified; in and de: proteins significantly increased or decreased respectively (p<0.05, Student’s *t* test). Total percentage of proteome altered in each comparison is indicated in purple. (**H-J**) pie charts indicate the significantly altered proteins assigned to regulation groups. Juvenile regulated proteins dominated by selective SD response, adults dominated by regulation from the wake/sleep cycle. (**K-M**) Venn diagrams depict overlap between ages in proteins significantly increased or decreased in each of the pairwise comparisons.

**Figure. 3.**
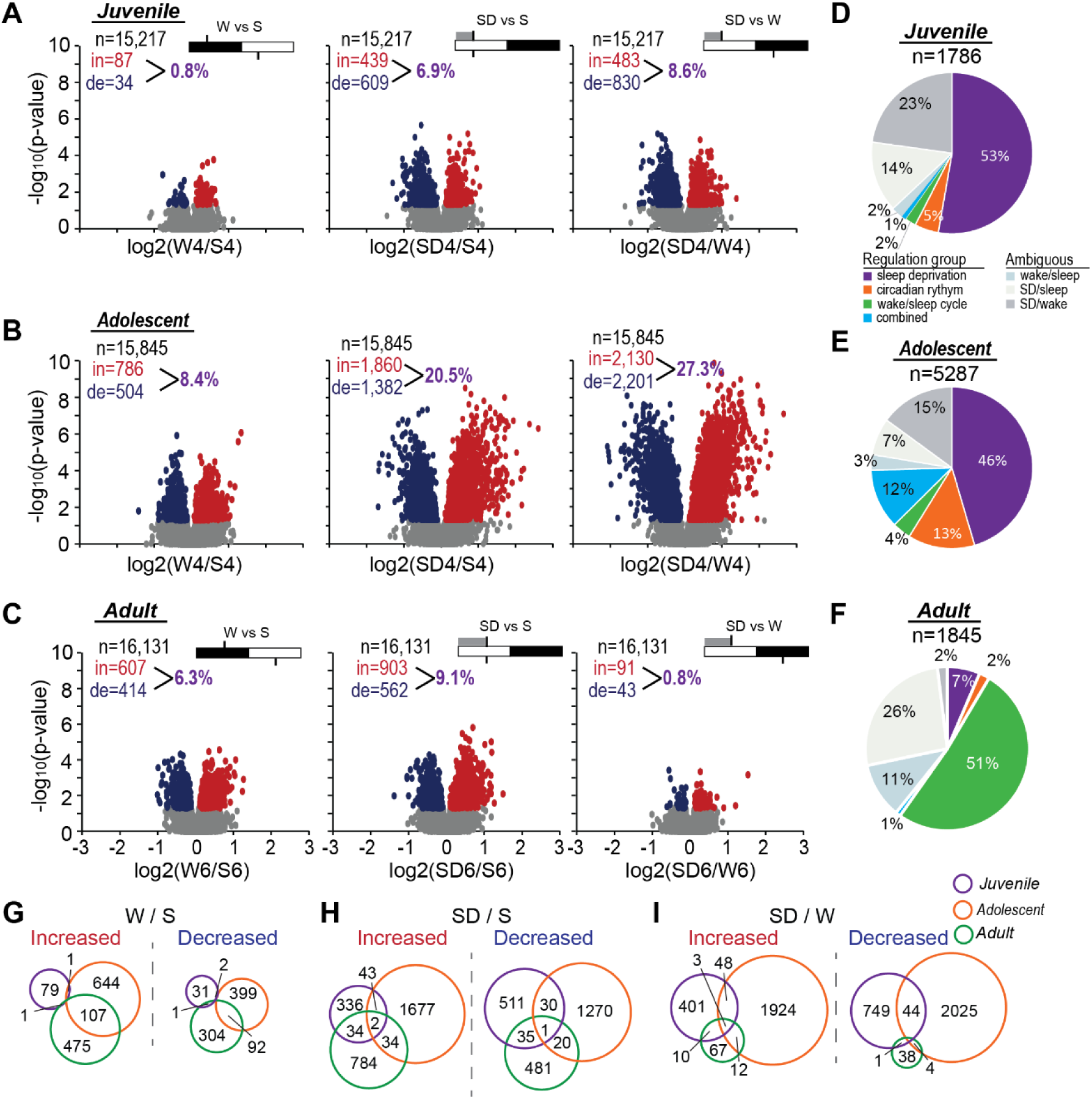
The developing synapse phosphoproteome is highly vulnerable to SD. (**A-C**) Volcano plots showing changes in synapse phosphopeptides in wake/sleep, SD/sleep, and SD/wake comparisons from juvenile, adolescent, or adult mice. N: number of phosphopeptides quantified; in and de: phosphopeptides significantly (p<0.05, Student’s *t* test) increased or decreased respectively. Total percentage of phosphoproteome altered in each comparison is indicated. (**D-F**) Pie charts indicate the significantly altered phosphopeptides assigned to regulation groups. Juvenile and adolescent regulated phosphopeptides dominated by selective SD response, adults dominated by regulation from the wake/sleep cycle. (**G-I**) Venn diagrams depict overlap between ages in phosphopeptides significantly increased or decreased in each of the pairwise comparisons. Minimal overlap between ages under all conditions examined.

By performing pairwise comparisons between wake/sleep/SD conditions, our experimental design allows for identification of synapse proteins regulated by the circadian rhythm (time of day), sleep-wake vigilance states, a selective response to SD, or a combination, in developing and adult mice, independent of sex (Fig. 2D and Fig. S2). We first identified significantly regulated synapse proteins/phosphopeptides (p<0.05) in one or more pairwise comparisons between conditions, followed by assignment to “regulation groups”: sleep/wake cycle, circadian rhythm, sleep deprivation, combined, or ambiguous (p<0.1 in 2 or 3 pairwise comparisons, described in Fig. S2). Note, biochemical analysis on bulk tissue is well suited to identify a unified, net effect of experimental conditions. In the juvenile W4/S4 comparison, we observed 55 significantly altered proteins, representing only 0.9% of the synapse proteome, indicating absence of an overt net biochemical change, consistent with reports that mixed types of plasticity co-occur during sleep at the juvenile age, including synapse strengthening and weakening/pruning^27,41^. In contrast, SD4/S4 and SD4/W4 comparisons showed 768 (13.0%) and 842 (14.3%) significantly altered proteins respectively (Fig. 2E), clearly indicating that SD drove a very large unified biochemical effect on juvenile synapses readily detected by our analysis.

Overall analysis of juveniles showed 1,152 proteins (19.5% of the total synapse proteome) significantly altered between the three conditions; of these a clear majority, 62%, showed a selective response to SD (Fig. 2H), compared to only ∼2% each for circadian rhythm or wake-sleep cycle, indicating the developing synapse proteome is profoundly sensitive to SD. Near identical conclusions are drawn when proteins are assigned to regulation groups using more stringent statistical cutoffs (Fig. S2).

A dramatic transition in synaptic responses occurs with maturation to adolescence and adulthood. Compared to the minimal W4/S4 changes seen in juveniles, we observe the emergence of a unified change between wake and sleep conditions where the W4/S4 comparison in adolescents revealed 351 (7.2%) proteins, and W6/S6 comparison in adults revealed 629 (9.5%) altered proteins (Fig. 2F and G). Overall, in adolescents we identified 796 (16.4%) regulated proteins between conditions. Interestingly, compared to juveniles, adolescents display a comparable reduction in the fraction of regulated proteins responding selectively to SD (22%), and emergence of a considerable component (34%) regulated by circadian rhythm, with only 3% selectively responding to wake-sleep cycle (Fig. 2I). Finally, in adults we identified 772 (11.7%) regulated proteins, of these the clear majority (68%) were regulated as a function of the wake-sleep cycle, with only 2-3% of proteins selectively responding to SD or circadian rhythm (Fig. 2J). Indeed, the SD6/W6 comparison in adults, where both conditions are awake but at opposite times of day, showed almost no differences. This latter point is entirely consistent with recent multi-omics studies examining the daily regulation of the adult forebrain synapse, reporting that the synaptic transcriptome (synapse localized mRNA) is regulated primarily as a function of the circadian rhythm, whereas the synaptic proteome and phosphoproteome are regulated almost exclusively as a function of wake-sleep states^9,10^. Consistent with adult resilience to SD4 in the NOR test (Fig. 1H), we find that SD4 treatment has a minimal effect on the adult synapse proteome compared to SD6 (Fig. S3A). Finally, comparison of significantly regulated proteins across ages and conditions revealed minimal overlap, indicating an almost entirely distinct proteomic response at each age, particularly notable for the juveniles (Fig. 2K-M). Gene ontology (GO) analysis further indicates distinct biological effects of SD across the ages tested here (Fig. S4).

### SD drives a predominant phosphorylation-based response after maturation to adolescence

Next, we examined the dynamics of the synaptic phosphoproteome in juveniles, adolescents, and adults. Previous phosphoproteomics studies revealed that the adult synapse phosphoproteome is regulated primarily as a function of the wake sleep cycle^10^ and that SD drives accumulation of brain wide protein phosphorylation, primarily on synaptic proteins^5^. Results of our analysis corroborate these prior findings from adults, but further indicate that the developing synapse phosphoproteome shows an entirely distinct SD response. Between comparisons we identified 1,786 regulated synaptic phosphopeptides from juveniles, representing 11.7% of the phosphoproteome, a majority of which showed a selective response to SD compared to only 5% responding to circadian rhythm (Fig. 3A and D), reminiscent of the SD-dominated proteome response. In adolescents we observed striking dynamics of the phosphoproteome in each comparison, far greater in magnitude than the other age groups. In total, 5,287 phosphopeptides (33.4%) were found to be significantly regulated, a large fraction of which (46%) showed a selective response to SD, and 13% were regulated by the circadian rhythm (Fig. 3B and E). In adults, SD6/W6/S6 analysis revealed 1,845 regulated phosphopeptides (11.4%), of which a majority (51%) showed a selective response to wake-sleep cycle, and only 7% showed a selective response to SD (Fig. 3C and F), corroborating prior literature^10^. As with the proteome, assignment of regulated phosphopeptides to regulation groups using a more stringent statistical cutoff resulted in near identical conclusions (Fig S2). Substantially fewer phosphorylation changes, 2.6% of the phosphoproteome, were observed in the adult SD4/W4/S4 analysis (Fig. S3B), consistent with a time of SD “dose response” of the phosphoproteome^5^. In agreement with conclusions from the synapse proteome analysis (Fig. 2), regulated phosphopeptides showed minimal overlap between ages and conditions, indicative of a distinct phosphorylation response to SD at each age (Fig. 3G-I). Accordingly, kinase substrate enrichment analysis (KSEA), which predicts kinase activity based on phosphorylation data^42^, suggests distinct patterns of kinase activation between ages (Fig. S5), Consistent with prior literature^5^, SD drove an asymmetric accumulation of phosphorylation in the SD/sleep comparison for adolescents and adults (Fig. 3B and C). Interestingly, this pattern was not observed in juveniles, which showed slightly more decreased phosphorylations (Fig. 3A). We additionally examined phosphorylation changes occurring on previously identified “sleep-need index phosphoproteins” (SNIPPs), mostly synaptic localized proteins demonstrated to preferentially accumulate phosphorylations in response to SD^5^.

Compared to sleep, SD drove an asymmetric increase in SNIPP phosphorylation in adolescents and adults, whereas juvenile mice displayed a largely symmetrical change in SNIPP phosphorylation (Fig. S6). This is again indicative of a molecularly distinct SD response in juveniles, and emergence of a phosphorylation-based response with maturation through adolescence. Notably, the synaptic phosphorylation response in adolescents was substantially greater compared to adults, affecting more than twice the number of phosphopeptides (Fig. 3B and E). Previous phosphoproteomic analysis indicated synapse phosphorylation accumulates in response to duration, “dosing”, of SD^5^. We speculate that exaggerated phosphorylation responses seen in adolescents may result from accelerated accumulation of SD-driven phosphorylation downstream of sleep promoting kinases^12–15^, perhaps due to blunted adolescent adaptations in response to SD^37^ (Fig. 1).

### Sleep deprivation in juveniles drives immediate impacts on key aspects of brain development and interacts with nodes of genetic risk for ASD

Recent mathematical modeling examining sleep across life stages and species predicted a unique developmental function of sleep in early life supporting brain growth, abruptly transitioning to a homeostatic function with maturation^33^. Accordingly, gene ontology (GO) and STRING pathway analysis of SD sensitive proteins revealed that SD in juveniles, but not adults, drove upregulation of dozens of synaptic proteins related to axon-growth, synaptogenesis, myelination, and formation of perineuronal nets (PNNs) (Fig. 4A, and Fig. S4). These pathways represent key aspects of brain development actively shaping neuronal networks during the juvenile period^43–48^. In addition, SD in juveniles but not older groups drove upregulation of synapse proteins related to excitatory synaptic strengthening and Hebbian LTP, including DLG4 (PSD95) and prominent excitatory synapse scaffolds, CaMKIIα, AMPA and NMDA-type glutamate receptors and auxiliary subunits (Fig. 4A, and Fig. S4)^36,49^. Accordingly, kinase activity prediction using KSEA indicates that SD in juveniles, and not older mice, drives activation of CaMKIIα/β/γ (Fig. 4B and fig. S5), Ca^++^ activated kinases well known for a prominent function in Hebbian LTP^49^. These results suggest that SD in juveniles drives heightened neuronal activity that aberrantly stimulates synapse growth and Hebbian potentiation, further supporting an important role of the wake sleep cycle in shaping the maturation of synapses during the juvenile period^27–29^.

**Figure. 4.**
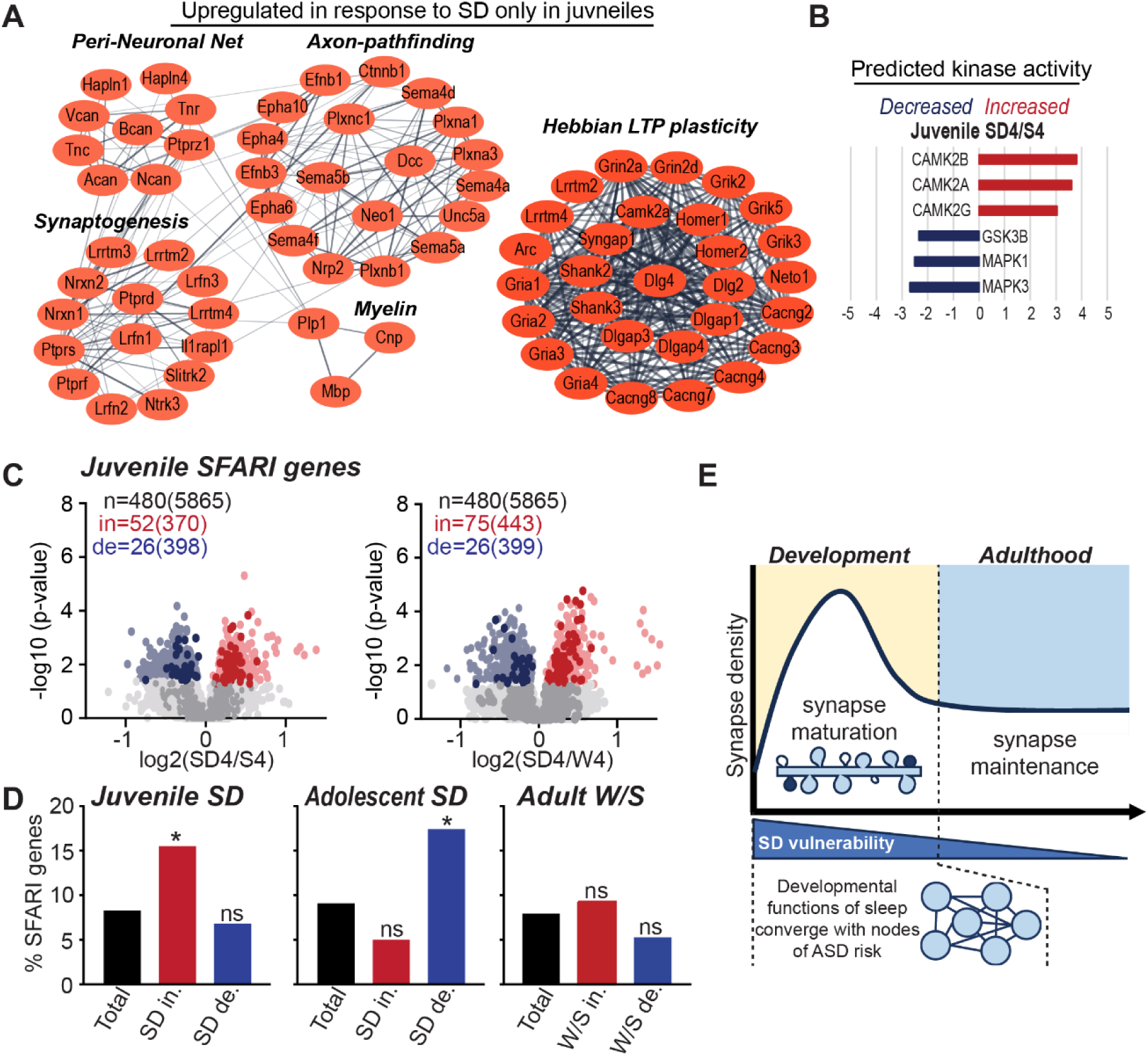
SD in juvenile impacts key aspects of synapse maturation, converging on nodes of ASD risk. **(A)** Synapse proteins upregulated in response to SD only in juveniles, related to brain development and synapse potentiation. Protein interaction network identified using gene ontology (GO) and STRING analysis. (**B**) KSEA indicates SD drives activation of CaMKII isoforms in juveniles. (**C**) Volcano plots: SD induced changes in SFARI risk genes (bold color). Number of SFARI genes detected (n), increased (in) or decreased (de) compared to total (in brackets) are indicated. (**D**) Bar charts, % of total or regulated proteins represented by SFARI genes: * indicates SD response is significantly enriched for SFARI genes compared to expected overlap from total, ns: not significant (hyper geometric test). Proteins upregulated by SD in juveniles or downregulated in adolescents are significantly enriched for SFARI genes. (**E**) Summary model.

Sleep disruption is a common and early phenotype associated with autism spectrum disorder (ASD), an early onset neurodevelopmental condition largely regarded as a synaptopathy^50^. In animal models, early life sleep disruption has been shown to drive lasting ASD-relevant changes in behavior^22–25^, interacting with underlying ASD genetic risk^24^. Of the hundreds of ASD risk genes now described^51^, compiled by SFARI (Simons Foundation Autism Research Initiative), a large fraction encode synaptic proteins. Together, these points led us to hypothesize that genetic risk for ASD and developmental sleep loss converge on maturation of the synapse. As expected, ASD risk genes are well represented in our synapse proteome datasets, representing ∼8% of detected synapse proteins. Importantly, we observed that proteins upregulated in response to SD in juveniles are significantly enriched (∼2 fold) for high-confidence ASD risk genes (Fig. 4C and D), supporting a sleep x gene interaction in ASD etiology^24^.

Adolescents showed a distinct pattern where SFARI genes are enriched in the proteins downregulated in response to SD (Fig. 4D). These findings suggest the development-specific functions of sleep that support synapse maturation converge on nodes of genetic risk for ASD in an age dependent manner (Fig. 4E). In contrast, synaptic proteins regulated by the wake/sleep cycle in adults (Fig. 2J) showed no enrichment for SFARI genes (Fig. 4D), further suggesting that as maturation progresses the vulnerability to SD decreases^24^ (Fig. 4E). Together, our behavioral, molecular, proteomic, and phosphoproteomic analyses (Fig. 1-3) all clearly indicate distinct responses to SD between juvenile, adolescent, and adult mice, indicative of unique functions of sleep, and associated vulnerabilities to SD across development.

## Discussion

Neuronal synapses have emerged as a distributed and conserved target for the restorative benefits of sleep and a locus of dysfunction in response to SD^4,5^. A prominent synapse-based model proposes information encoding during wake drives synaptic potentiation, perturbing synapse homeostasis, which is restored during sleep by broad synapse weakening through homeostatic scaling-down^6^. This restorative plasticity involves broad dephosphorylation of AMPA-type glutamate receptors, and signaling through metabotropic glutamate receptors mGluR1/5 activated by homeostatic scaling factor *Homer1a*^7,35,36^, molecules heavily implicated in sleep homeostasis^38,52^. Recent multi-omics efforts expand this view, showing the adult forebrain synapse proteome and phosphoproteome undergo profound remodeling, involving hundreds of molecules, driven almost entirely by wake sleep states^9,10^; our current results are entirely consistent and corroborate these findings in adults, whereas results from developmental time points indicate sleep exerts distinct synaptic functions, consistent with recent mathematical modeling^33^.

Contrasting reports between developmental and adult time points has driven controversy on the synaptic basis of sleep function^26^. Indeed, very few studies have engaged in systematic analysis of sleep function or response to SD across developmental and adult time points^37,53^. Forebrain synapses undergo well characterized dynamics during postnatal development, beginning with a period of synaptogenesis, followed by experience-dependent pruning^17^. During the transition to adulthood, overall synapse density stabilizes, the turnover of individual synapses reduces dramatically, and many synapses are maintained for long periods, perhaps the remainder of the lifespan^54,55^. Therefore, we speculate sleep engages distinct plasticity types during the period of developmental dynamics and adult maintenance (Fig. 4E). In adults, homeostatic scaling-down acts on pre-existing synapses to renew daily learning capacity. In contrast, during juvenile development, waking experience drives synaptogenesis, while sleep is critical for the selection of synapses to be pruned and for the maturation and growth of the remaining synapses^27–30^, facilitating circuit refinement. Acute SD in juveniles drives changes in the synapse proteome consistent with activation of Hebbian LTP, stimulation of synaptogenesis, and deposition of peri-neuronal nets (Fig. 4A). These findings support the view that ongoing synapse maturation is regulated by the wake/sleep cycle and inform possible mechanisms by which sustained developmental sleep disruption, commonly reported in ASD, drives long lasting effects on brain function and behavior^22–25^. Indeed, our analysis shows that juvenile vulnerability to SD and ASD genetic risk converge on synapse development. Finally, our findings, together with a prior systematic EEG analysis show that juveniles lack well characterized adaptations to SD: increased NREM-SWA^37^, dark phase sleep rebound, induction of *Homer1a*, and accumulation of synaptic protein phosphorylation, potentially exposing the developing brain to greater vulnerabilities of SD. A recent RNAseq analysis from frontal cortex also shows almost no overlap in SD-induced gene expression changes between juvenile and adult mice^53^. Taken together, our findings suggest sleep plays a distinct and important role supporting brain maturation and highlight the developing synapse as a major node of vulnerability to sleep loss relevant for childhood onset forms of neurodevelopmental disorders.

Computer modeling^33^ and our new findings indicate very different response to sleep or SD between juvenile and adult time points (Fig. 4E). Our systematic analysis identified that an important developmental transition occurs between the ages of P35 and P42 (Fig. 1), notably a period marking the end of critical period plasticity, stabilization of forebrain synapse density, and the emergence of sexual maturity in mice^17,56^. At this age, here referred to as adolescence, we see emergence of adult-like broad dephosphorylation of GluA1 AMPARs, consistent with a transition in sleep function towards synapse homeostasis and maintenance^7^. This period also marks the emergence of adult-like responses to SD, albeit blunted, including dark phase rebound, induction of *Homer1a* (Fig.1), and increased NREM-SWA^37^. However, proteomic and phosphoproteomic analysis of forebrain synapses (Figs. 2 and 3) further identify the adolescent age as a unique epoch of development, distinct from adulthood. Where the adult synapse is regulated almost exclusively as a function of wake/sleep states^9,10^, the adolescent synapse proteome and phosphoproteome exhibit sensitivity to SD and a notable component regulated by the circadian rhythm (time of day). This circadian component is not seen in juvenile mice.

Therefore, we speculate that as synapse densities stabilize and adult-like homeostatic plasticity becomes prominent, the circadian rhythm instructs the daily remodeling of the synapse, which subsequently becomes dominated by wake/sleep states with further maturation to adulthood. Just as the juvenile age exhibits distinct vulnerabilities to SD relevant in early onset neurodevelopmental conditions such as ASD^18^, this transitional adolescent age is likely associated with distinct vulnerabilities to sleep and circadian disruption. This may be relevant for adolescent onset neuropsychiatric illnesses such as schizophrenia, where sleep and circadian disruption are prominent comorbidities^19^. Our findings support that sleep function undergoes important transitions across development, offering some resolution to controversies in this field. Recognition of the unique functions of sleep across development will lead to further understanding of the role of sleep in lifelong brain health.

## Materials and Methods

### Mice

All animal procedures were approved by the Institutional Animal Care and Use Committee of the University of North Carolina (UNC) at Chapel Hill and were performed in accordance with the guidelines of the U.S. National Institutes of Health. All experiments were performed using C57Bl/6J mice of both sexes purchased from Jackson Labs, or bred in house. Adult mice for all experiments were purchased from Jackson Labs and acclimated to UNC housing for at least 2 weeks prior to the start of the experiment. For experiments using juvenile or adolescent mice, experimental animals were bred in house with adult breeders purchased from Jackson Labs. Breeders for our colony were replaced every 3 months with mice supplied from Jackson Labs. Target experimental ages for adolescents was P42 and adults P90. For juveniles, target age for biochemistry experiments was P21, target age for behavior studies was P28. Specific age ranges are stated below for each experiment.

### Sleep recordings and calculations

Sleep-wake behavior of juvenile (postnatal day, p25-33), adolescent (p41-45), and adult (p70-100) C57BL/6J mice was recorded continuously using a non-invasive piezoelectric monitoring system (PiezoSleep 2.0; Signal Solution, LLC, Lexington, KY). Separate cohorts were generated for each of the age groups. First, mice were transferred to our satellite facility, maintained on a 12hr:12hr light:dark cycle. Mice were housed individually (15.5 cm^2^ cages) and provided with bedding, water, and food. Following a 36-48hr acclimation period, sleep and wake behavior was recorded for at least 24 hours to establish a baseline. Next, mice were subjected to a 4hr gentle handling sleep deprivation (SD4) at the onset of the light cycle (zeitgeber time, ZT0-4) and allowed to recover for the remainder of the day (ZT4 – ZT24). During SD, wakefulness was assessed by continued observation from experimenters as described previously ^39^.

The PiezoSleep monitoring system utilizes a piezoelectric mat positioned underneath the mouse cage to detect vibrational movements, such as breathing. A customized statistics software (SleepStats, Signal Solution, Lexington, KY) processes these signals through an algorithm, distinguishing distinct respiratory patterns associated with sleep and wake states. Sleep is typified by periodic (2-3 Hz), regular amplitude signal, a common respiratory feature during mouse sleep states. In contrast, wakefulness is identified by the absence of standard sleep signals and the presence of irregular, higher amplitude signals, typically observed during voluntary movement (including subtle movements seen in quite wake). Using a linear discriminant classifier algorithm, the piezoelectric signals were classified as “sleep” or “wake” in 2-second epochs. Sleep-wake thresholds were automatically calculated for each individual mouse. To exclude brief or ambiguous arousals, a sleep bout is initiated when a 30 sec epoch contains greater than 50% sleep and is terminated when a 30 sec epoch contains less than 50% sleep. Validation of this software has been conducted using a combination of electroencephalography, electromyography, and visual assessment ^57–59^.

Sleep amount generated through the SleepStats software was analyzed to characterize sleep-wake patterns on the day of sleep deprivation (SD) and the day prior (Baseline). Sleep amount data was binned into 1hr intervals to generate daily sleep traces and calculate average sleep amount during the light and dark cycles. Sleep amount lost during SD (SD deficit) was calculated by subtracting baseline sleep amount from SD sleep amount during ZT0-ZT4. Sleep amount recovered was calculated separately for the light and dark cycles, by subtracting baseline sleep amount from SD sleep amount during ZT4-ZT12 (light rebound) and ZT12-ZT24 (dark rebound), respectively. Total sleep recovery was then calculated as a percentage of the average SD deficit of each age group. A 2-way ANOVA followed by a Sidak’s multiple comparisons test was used to analyze total sleep time. Sleep recovery metrics were analyzed with a one-way ANOVA followed by a Tukey’s multiple comparisons test.

### Quantitative Real-Time PCR

Forebrain (cortex and hippocampus) were dissected from juvenile (P21-22), adolescent (P42-46), and adult (P90) mice. (mRNA was purified using the RNeasy Plus Mini Kit (Qiagen 74136) according to the manufacturer’s instruction. RNA was reverse transcribed into cDNA using the High-Capacity cDNA Reverse Transcription Kit (Applied Biosystems 4374966). Quantitative real-time PCR (RT-qPCR) was performed using the QuantStudio 7 Flex (Applied Biosystems) using Taqman Fast Advanced Master Mix (Applied Biosystems 4444557) and Taqman primer/probes for GAPDH (Invitrogen, Mm99999915_g1), Per2 (Invitrogen, Mm00478099) and Homer1a (Invitrogen custom Taqman Assay, APNKUGG). Expression data were calculated as fold gene expression using 2^-ΔΔCt^ method with GAPDH as the reference gene and normalized to the sleep (ZT4) condition. Four independent technical replicates were performed in each individual experiment. The resulting data was analyzed with a 2-way ANOVA followed by Tukey’s multiple comparisons test.

### Novel Object Recognition (NOR) assay and analysis

Mice were tested at one of three age points: juvenile (P28); adolescent (P42); and adult (P100). All experimental manipulations took place in a lab-maintained behavior assay room (“behavior room”). The experimental apparatus was composed of four open-topped cubes arranged in a quadrant formation at table height. The cubes were 40cm3 and were lined with napped gray Kydex. One wall in each cage had a visual cue so that animals could spatially orient themselves. Small action cameras (AKASO EK7000 Pro) were positioned directly over each chamber to record the mice during the experiment. Each chamber also had a dedicated light source directly overhead to allow for dim lighting conditions, with the overhead lights off, during the NOR testing. The objects were pairs of identical glass salt shakers or similarly sized black plastic canisters. The objects were weighted to prevent tipping or sliding, and they were positioned within each cage using a measured placement guide. Chambers and objects were cleaned with ethanol between subjects, and subjects were returned to the same chamber for training and testing.

For three days prior to training, all mice were lightly handled for 10 minutes to acclimatize the mice to the experimenters. By the third day, all animals could perch calmly in the experimenter’s palm without voiding or trying to escape.

On training day, all subjects in a cohort were brought to the behavior room in their home cages at ZT1. Each mouse freely explored one chamber with two identical objects in fixed locations for 10 minutes. Four animals were trained simultaneously. After 10 minutes, mice were returned to their home cages with their cage mates and either allowed to have undisturbed sleep (control) or gentle handling sleep deprivation for 4 hours (SD4) or 6 hours (SD6). Sleep deprivation took place in the home cage in the behavior room.

On testing day, the following morning, all subjects were brought back to the behavior room in their home cages at ZT2. Each mouse was returned to its training cage and allowed to freely explore for 10 minutes one familiar training object and one novel object (whichever class of object was not used for that mouse’s training) in the same fixed locations as during training. Four animals were again tested simultaneously. The experiment was concluded once each mouse completed its testing exposure, and the chambers and objects were deep cleaned and sanitized.

#### Scoring

A scoring function was developed in-house using Oracle OpenJDK Version 17.0.1. It was used in the IntelliJ IDEA 2021.3.1 IDE by scorers to record beginning and end points, in milliseconds, of an interaction with each object, while viewing the video recording of the testing sessions. The scorers were blind to mouse sleep condition and familiar/novel object status during the time of scoring. The function also recorded all beginning points of interactions (“Time stamps”), as well as the beginning and end time points of the scoring process/video. Interactions were defined as any continuous period during which the mouse was within one head-length from the object of interest and looking directly at the object. If the mouse looked away or climbed on top of the object, the interaction period ended.

#### Data analysis

Values for total time spent interacting with each object (Total Interaction Time) and total novel object interaction time divided by total overall object interaction time (Discrimination Index) were calculated and subsequently exported to GraphPad Prism 9.5.1 for figure development and statistical analyses. Statistical values for Total Interaction Time were calculated using paired t-tests, comparing within-mouse novel object and familiar object interaction time values. Statistical values for Discrimination Index were calculated using unpaired t-tests comparing control and sleep-deprived mice within each age group.

### Mouse collection and PSD preparation

Developing mice (P21-P49) were sacrificed at ZT4 (sleep), ZT16 (wake), or at ZT4 following SD4. Adult mice (P70-100) were sacrificed undisturbed at ZT4/ZT16 or immediately following SD4 (ZT4), or undisturbed at ZT6 (sleep), ZT18 (wake), or immediately following SD6 (ZT6). An equal amount of male and females were used for each condition and age, allowing us to determine changes independent of sex. Once sacrificed, the forebrain was dissected and immediately flash frozen.

Frozen forebrains were homogenized using 15 strokes from an ice-cold homogenizer in ice cold homogenization solution (320mM sucrose, 10mM HEPES pH 7.4, 1mM EDTA, 5mM Na pyrophosphate, 1mM Na_3_VO_4_, 200nM okadaic acid, protease inhibitor cocktail). The brain homogenate was then spun at 1,000xg for 10min at 4°C to obtain the P1 (nuclear) and S1 (post-nuclear) fractions. The S1 fraction was layered on top of a discontinuous sucrose density gradient (0.8M, 1.0M or 1.2M sucrose in 10mM HEPES pH 7.4, 1mM EDTA, 5mM Na pyrophosphate, 1mM Na3VO4, 200nM okadaic acid, protease inhibitor cocktail) and subjected to ultra-centrifugation at 82,500xg for 1.5hr at 4°C. The brain material present at the interface of the 1.0M and 1.2M sucrose buffers (synaptosomes) was collected. Synaptosomes were diluted using 10mM HEPES pH7.4 (containing protease and phosphatase inhibitors) to 320mM sucrose concentration. The diluted synaptosomes were pelleted by centrifugation at 100,000xg at 4°C for 20min. The pellet was resuspended in 50mM HEPES pH 7.4 and then mixed with an equal part 1% Triton X-100 (both solutions contained protease and phosphatase inhibitors). The mixture was incubated at 4°C for 15 minutes before being pelleted by centrifugation at 32,000xg for 20min at 4°C to yield the post-synaptic density (PSD) fraction. The pellet was then resuspended in 50mM pH 7.4 (plus protease and phosphatase inhibitors). The protein concentration was determined using Bradford assay. For proteomics and phophosproteomics 4 mice were pooled together for each individual sample analyzed. For western blots, individual mice were processed and analyzed.

### Antibodies

The following mouse primary antibodies were used for western blotting: GluA1 (neuromab 75-327, 1:1000) and PSD-95 (Neuromab 75–028, 1:1,000,000).

The following rabbit primary antibodies were used: GluA1 phospho S845 specific (Cell Signaling AB5849, 1:1000) and GluA1 phospho specific (Millipore AB5847, 1:1000)

### Proteomics and Phosphoproteomic preparation and analysis

A total of four sample sets (juvenile, adolescent, adult SD4/ZT4/ZT16, and adult SD6/ZT6/ZT18) were processed for proteomics and phosphoproteomics analysis. PSD lysates (0.5 mg per sample) were reduced with 5mM DTT for 45 min at 37°C, alkylated with 15mM iodoacetamide for 30 min in the dark at room temperature, then diluted to 1M urea with 50mM ammonium bicarbonate (pH 7.8). Samples were digested with LysC (Wako, 1:50 w/w) for 2 hr at 37°C, then digested with trypsin (Promega, 1:50 w/w) overnight at 37°C. The resulting peptide samples were acidified, desalted using desalting spin columns (Thermo), then the eluates were dried via vacuum centrifugation. Peptide concentration was determined using Quantitative Colorimetric Peptide Assay (Pierce). Twelve total samples and four pooled samples per experiment were labeled with TMTpro (Thermo Fisher) for 1 hr at room temperature. Prior to quenching, the labeling efficiency was evaluated by LC-MS/MS analysis. After confirming >98% labeling efficiency, samples were quenched with 50% hydroxylamine to a final concentration of 0.4%.

Labeled peptide samples were combined 1:1, desalted using Thermo desalting spin column, and dried via vacuum centrifugation. The dried TMT-labeled samples (four different set in total) were fractionated using high pH reversed phase HPLC ^60^. Briefly, the samples were offline fractionated over a 90 min run, into 96 fractions by high pH reverse-phase HPLC (Agilent 1260) using an Agilent Zorbax 300 Extend-C18 column (3.5-µm, 4.6 × 250 mm) with mobile phase A containing 4.5 mM ammonium formate (pH 10) in 2% (vol/vol) LC-MS grade acetonitrile, and mobile phase B containing 4.5 mM ammonium formate (pH 10) in 90% (vol/vol) LC-MS grade acetonitrile. The 96 resulting fractions were then concatenated in a non-continuous manner into 24 fractions and 5% of each were aliquoted, dried down via vacuum centrifugation and stored at −80°C until further analysis. The remaining 95% of each fraction was further concatenated into 6 fractions and dried down via vacuum centrifugation. For each fraction, phosphopeptides were enriched using the High Select Fe-NTA kit (Thermo) per manufacturer protocol, then the Fe-NTA eluates were dried down via vacuum centrifugation and stored at −80°C until further analysis.

#### LC/MS/MS Analysis

Each experiment consisting of 24 fractions for the proteome analysis and 6 fractions for the phosphoproteome analysis were analyzed by LC/MS/MS using an Easy nLC 1200-Orbitrap Fusion Lumos Tribrid or an Ultimate 3000-Exploris480 (Thermo Scientific). On the Lumos platform, samples were injected onto an Easy Spray PepMap C18 column (75 μm id × 25 cm, 2 μm particle size) (Thermo Scientific) and separated over a 120 min method. The gradient for separation consisted of 5–42% mobile phase B at a 250 nl/min flow rate, where mobile phase A was 0.1% formic acid in water and mobile phase B consisted of 0.1% formic acid in 80% acetonitrile. For the proteome fractions, the Lumos was operated in SPS-MS3 mode ^61^, with a 3s cycle time. Resolution for the precursor scan (m/z 400–1500) was set to 120,000 with a AGC target set to standard and a maximum injection time of 50 ms. MS2 scans consisted of CID normalized collision energy (NCE) 32; AGC target set to standard; maximum injection time of 50 ms; isolation window of 0.7 Da. Following MS2 acquisition, MS3 spectra were collected in SPS mode (10 scans per outcome); HCD set to 55; resolution set to 50,000; scan range set to 100-500; AGC target set to 200% with a 100 ms maximum inject time. For the phosphoproteome fractions, which were analyzed in duplicate, the Lumos was operated in MS2 mode ^62,63^ with a 3s cycle time. Resolution for the precursor scan (m/z 400–1500) was set to 60,000 with a AGC target set to standard and a maximum injection time of 50 ms. For MS2 scans, HCD was set to 35; AGC target set to 200%; maximum injection time of 120 ms; isolation window of 0.7 Da; resolution set to 50,000. On the Exploris platform, the proteome and phosphoproteome fractions were analyzed using a turboTMT method ^61^. Samples were injected onto an Ion Opticks Aurora C18 column (75 μm id × 15 cm, 1.6 μm particle size) and separated over a 70 or 100 min method. The gradient for separation consisted of 5–42% mobile phase B at a 250 nl/min flow rate, where mobile phase A was 0.1% formic acid in water and mobile phase B consisted of 0.1% formic acid in 80% ACN. The Exploris480 was operated in turboTMTpro mode with a cycle time of 3s. Resolution for the precursor scan (m/z 375–1400) was set to 60,000 with a AGC target set to standard and a maximum injection time set to auto. MS2 scans (30,000 resolution) consisted of higher collision dissociate (HCD) set to 38; AGC target set to 300%; maximum injection time set to auto; isolation window of 0.7 Da; fixed first mass of 110 m/z.

#### Data Analysis

Raw data files were processed using Proteome Discoverer version 2.5, set to ‘reporter ion MS2’ or ‘reporter ion MS3’ with ‘16pex TMT’. Peak lists were searched against a reviewed Uniprot mouse database (downloaded Feb 2021 containing 17,051 sequences), appended with a common contaminants database, using Sequest HT within Proteome Discoverer. Data were searched with up to two missed trypsin cleavage sites, fixed modifications: TMT16plex peptide N-terminus and Lys, carbamidomethylation Cys, dynamic modification: N-terminal protein acetyl, oxidation Met. For phosphoproteome data, additional dynamic modification: phosphorylation Ser, Thr, Tyr. Precursor mass tolerance of 10ppm and fragment mass tolerance of 0.02 Da (MS2) or 0.5 Da (MS3). Peptide false discovery rate was set to 1%.

Reporter abundance was calculated based on intensity, and for MS2 data, co-isolation threshold was set to 50; for MS3 data, SPS mass matches threshold was set to 50 and co-isolation threshold was set to 100. For proteome data, razor and unique peptides were used for quantitation. Proteins and phosphopeptides with >50% missing TMT intensities across samples were removed. Student’s t-tests were conducted within Proteome Discoverer, and a p-value <0.05 was considered significant. Log2 fold change ratios were calculated for each pairwise comparison. For the proteomic data set, proteins with PSM = 1 & only one unique peptidewere removed. Proteins identified as contaminants were removed. For the phosphoproteomic dataset, only phosphopeptides identified every sample for a given age group were analyzed.

### Bioinformatics and analysis

#### GO term and STRING analysis

To determine relevant biological and mechanistic pathways effected by sleep or sleep deprivation both the Gene Ontology data base and STRING analysis were used in conjunction with literature review. Significantly changing protein gene names were input into the Gene Ontology (GO) database ^64,65^. The GO database utilizes the PANTHER data base classification system to indicate groups of proteins that are represented significantly more than expected and identifies the associated GO terms ^66^. A false discovery rate (FDR) of 0.05 was the cut off for determining GO terms. GO terms were prioritized based on FDR, the PANTHER data base classification of GO term families, and with manual curation, using GO term curation guidelines ^67^. STRING analysis was performed on high confidence significantly changing proteins ^68^ on Cytoscape ^69,70^. A confidence score of 0.4 with no maximum interactors were used as cut offs. Apparent clusters were chosen and identified with GO term analysis and manual identification.

#### KSEA

Kinase-Substrate Enrichment Analysis (KSEA) was used to determine the predicted kinase activity from the phosphoprotoemic data set ^71–73^. The significantly changing phosphopeptides for each comparison were input in to the KSEA app software (4). Both the PhosphoSitePlus and NetworKIN databases were used. A score cutoff of 2 was used for the NetworKIN results. The kinases shown were predicted to significantly change in activity (p<0.05) and have 3 or more substrates.

## Acknowledgments

The authors would like to thank Lisa Sharek for technical assistance.

## Funding

Simons Foundation Autism Research Initiative #970806 and #385188 (GHD) This research is based in part upon work conducted using the UNC Proteomics Core Facility, which is supported in part by NCI Center Core Support Grant (2P30CA016086-45) to the UNC Lineberger Comprehensive Cancer Center.

---

**Figure S1.**
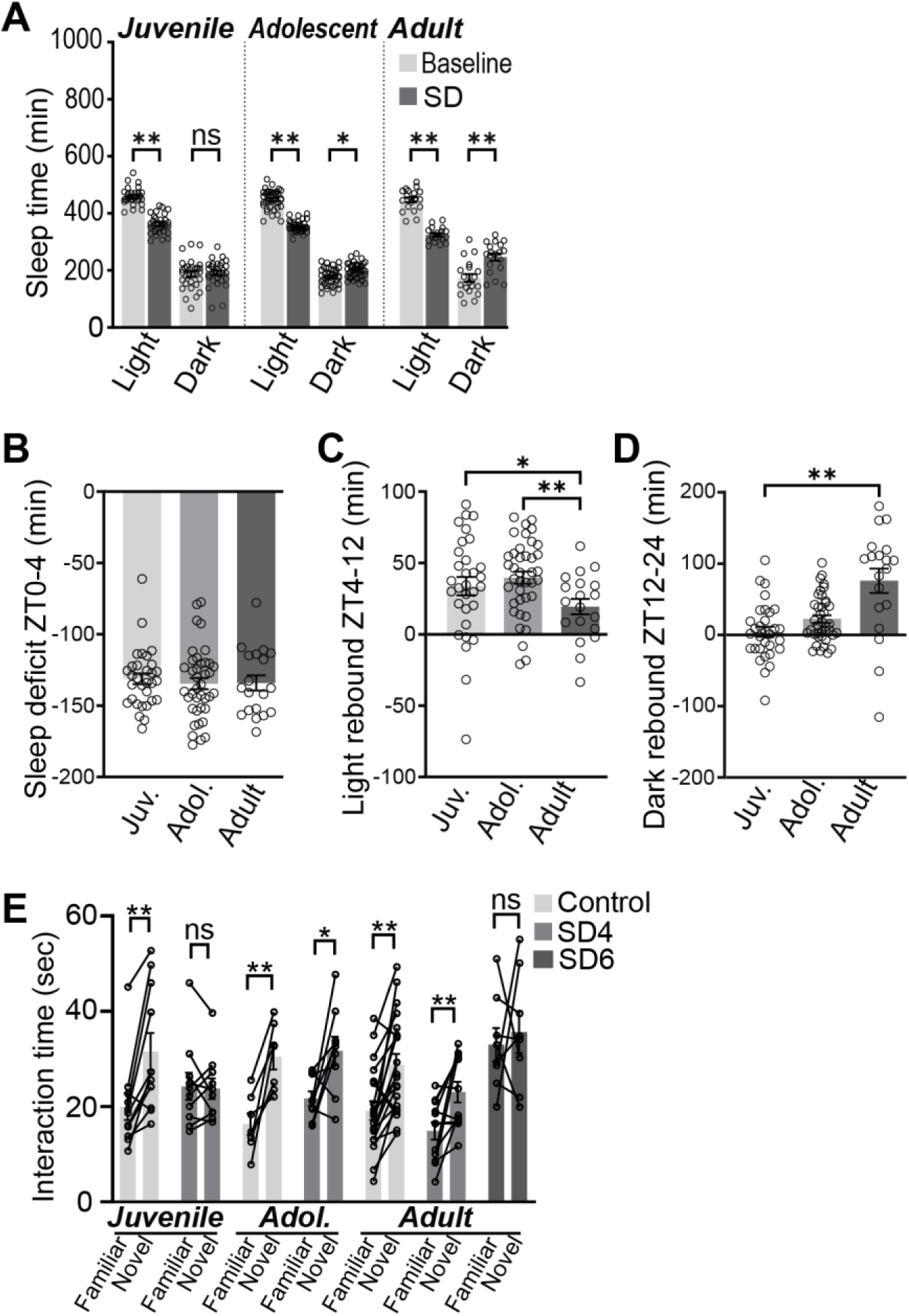
Differential sleep rebound responses across ages and effects of SD on NOR task interaction times. (**A**) Total sleep time in the light and dark phase during baseline or SD+recovery. Significant dark phase sleep rebound is measured in adolescents and adults, but not juveniles. *P<0.05, **P<0.01, ns: not significant, 2-way ANOVA, Sidak’s correction. (**B**) Sleep deficit calculated from difference in sleep amount from ZT0-4 between baseline and SD day. (**C and D**) comparison of light phase (**C**) and dark phase (**D**) sleep rebound between ages. Rebound measured as increased light phase ZT4-12, or dark phase (ZT12-24) sleep amount during SD+recovery over baseline. *P<0.05, **P<0.01, 1-way ANOVA, Tukey’s correction. (**F**) Interaction time with familiar and novel objects in the novel object recognition task. *P<0.05, **P<0.01, ns: not significant, paired Student’s t-test.

**Figure S2.**
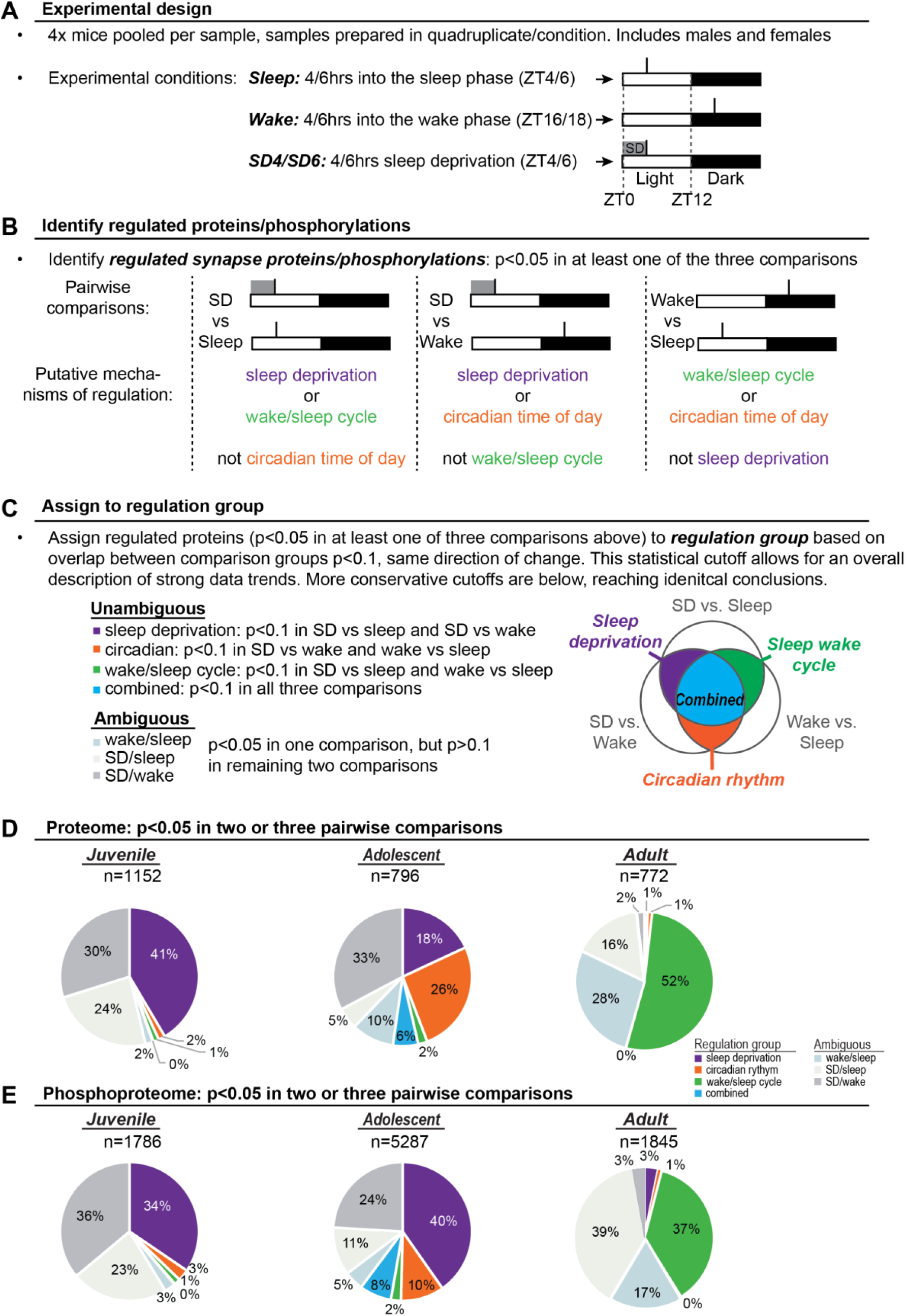
Interpretation of significantly regulated proteins and phosphopeptides. **(A)** experimental design. Mouse forebrain samples collected during wake or sleep phases, or following SD, 4 mice pooled per sample, samples prepared in quadruplicate. (**B**) pairwise comparisons between conditions identifies significantly regulated proteins. (**C**) regulated proteins assigned to regulation groups based on p-value cutoff. (**D and E**) Assignment of proteins (D) or phosphopeptides (E) to regulation groups based on more conservative statistical cutoffs: p<0.05 in two or three of the pairwise comparisons.

**Fig. S3.**
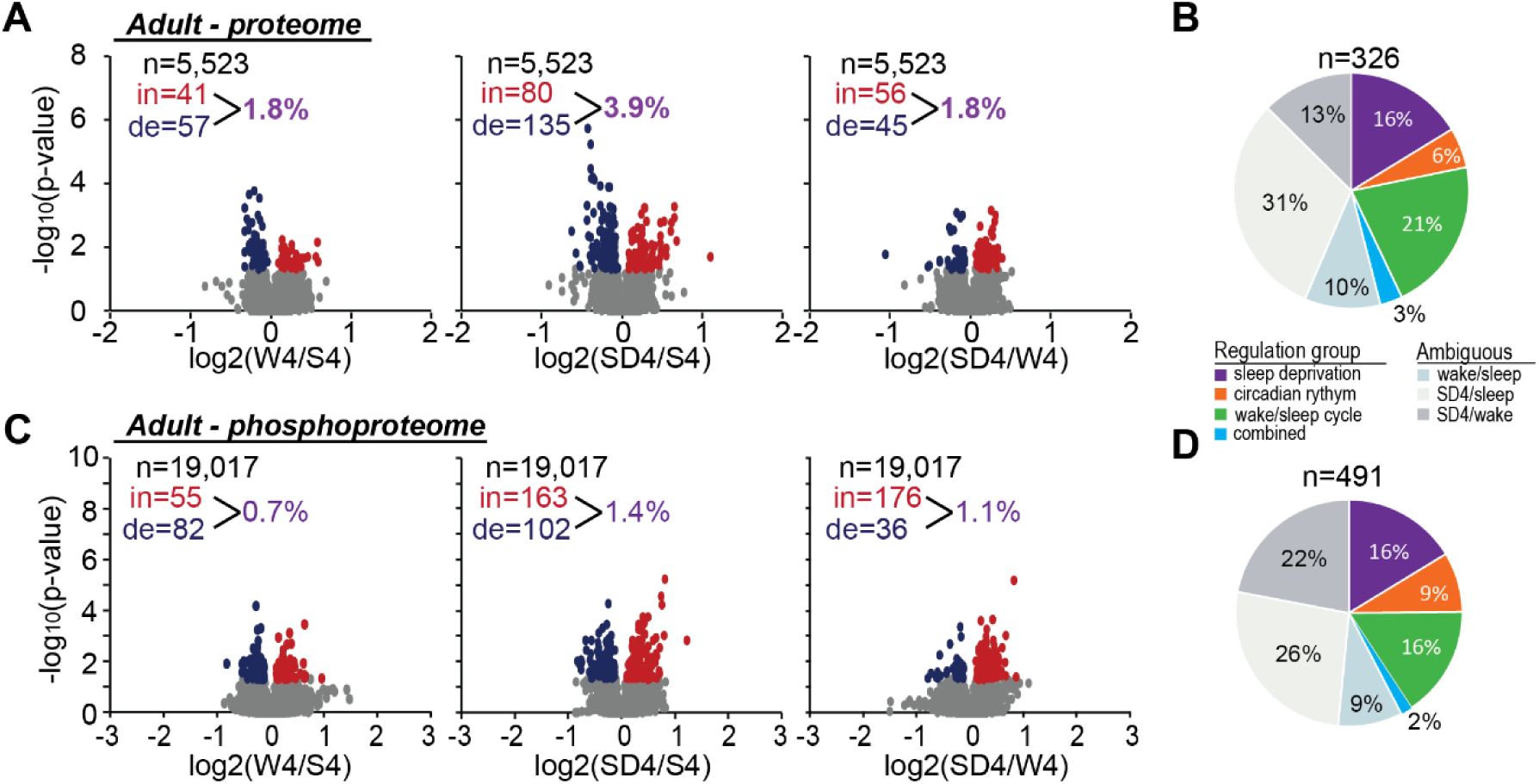
SD4 effects on adult synapse proteome and phosphoproteome. (**A**) volcano plots showing changes in synapse proteins in W4/S4, SD4/S4, and SD4/W4 comparisons from adult mice. N: number of proteins quantified; in and de: proteins significantly (p<0.05, Student’s *t* test) increased or decreased respectively. Total percentage of proteome altered in each comparison is indicated. (**B**) pie charts indicate the significantly altered proteins assigned to regulation groups. (**C and D**) Volcano plots and pie charts of adult phosphoproteome from W4/S4/SD4 conditions.

**Fig. S4.**
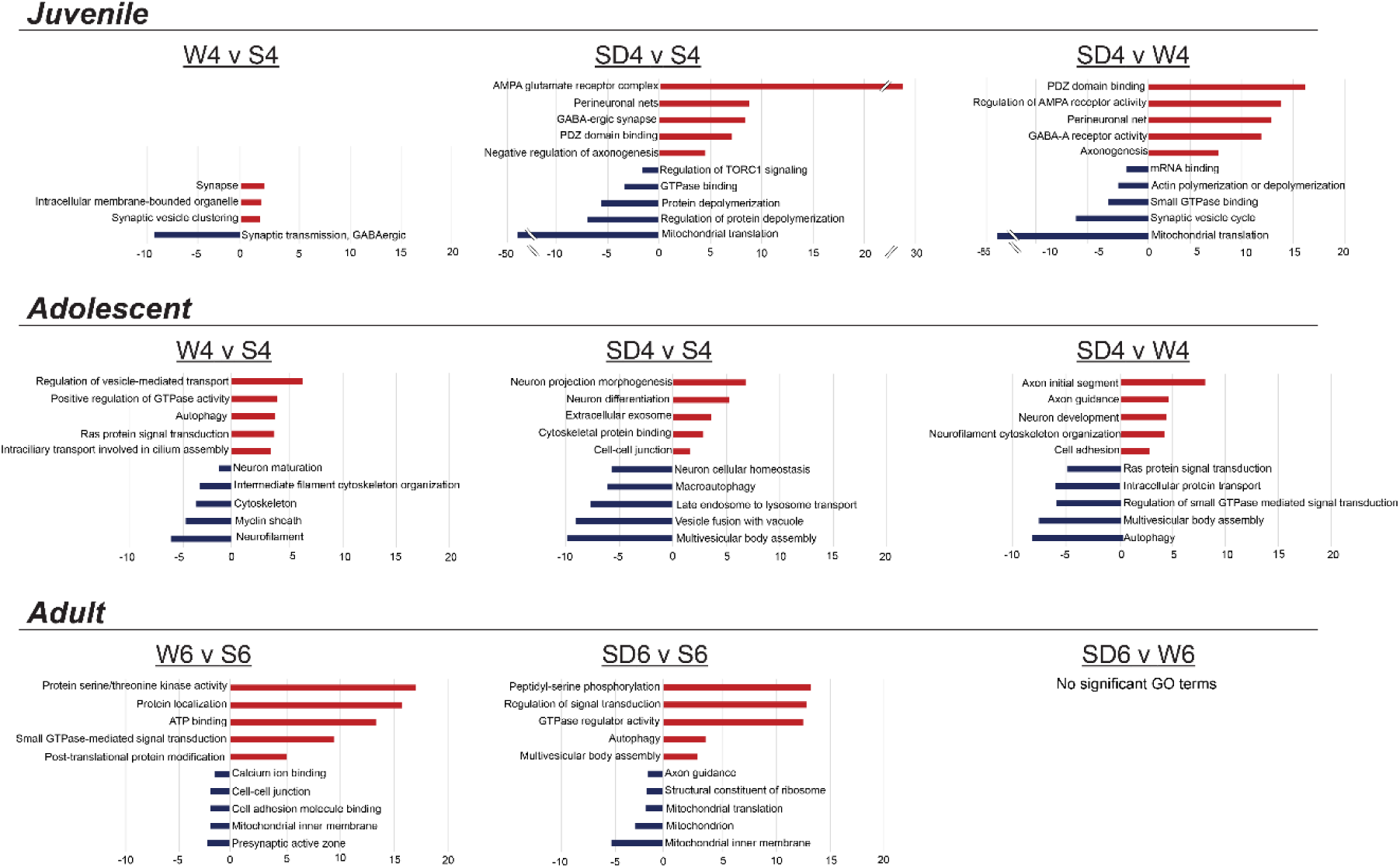
GO terms describing wake, sleep, and SD proteomic comparisons across ages. Gene ontology (GO) terms associated with synapse proteome analysis conducted from juvenile, adolescent, and adult mice. Depicted based on pairwise comparisons between wake, sleep, and SD conditions.

**Fig. S5.**
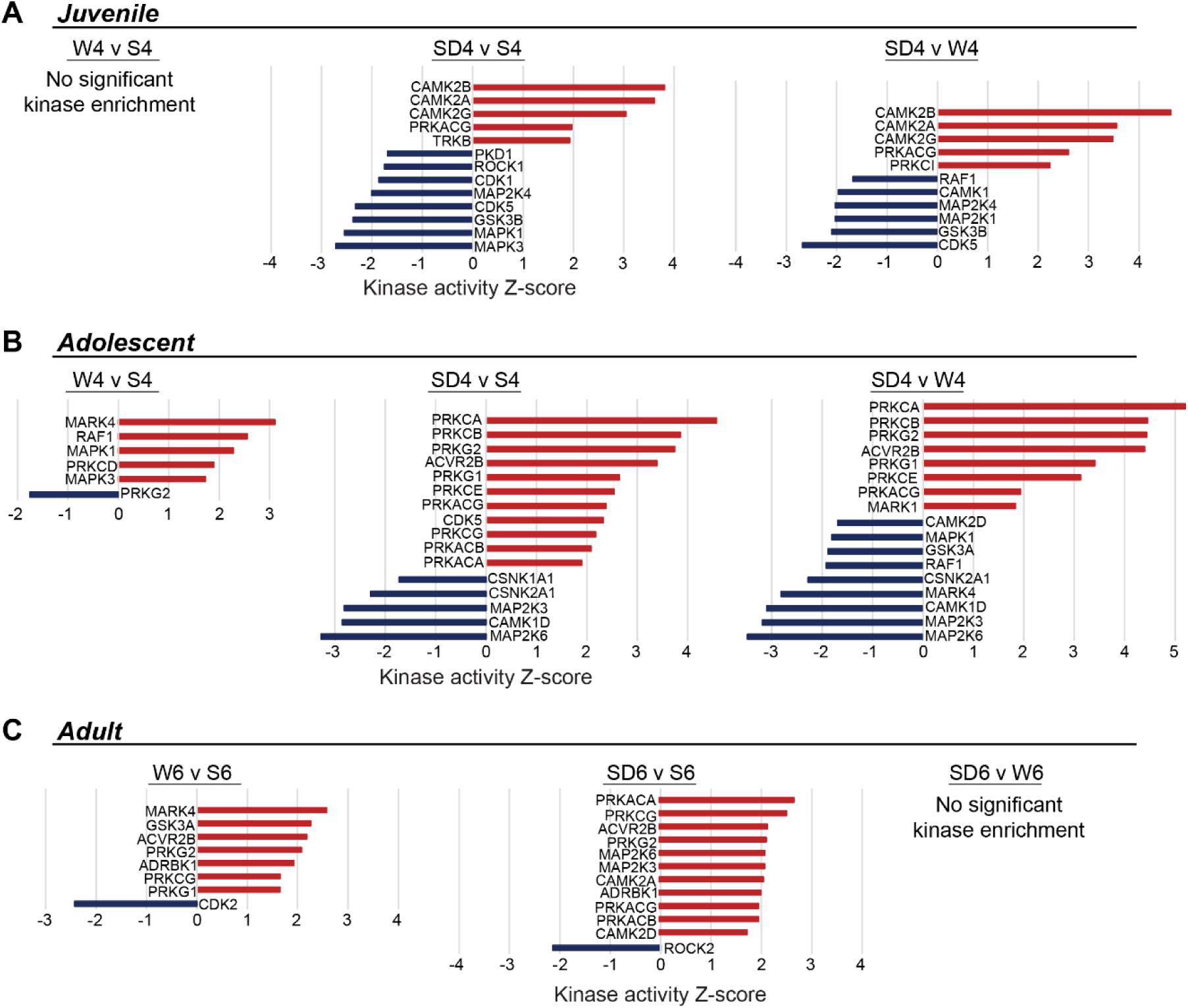
Kinase activity predictions from developing and adult synapse phosphoproteome. (**A-C**) Kinase substrate enrichment analysis (KSEA) was used to predict upregulated (red) or downregulated (blue) kinase activity in WvS, SDvS, and SDvW comparisons from each age.

**Fig. S6.**
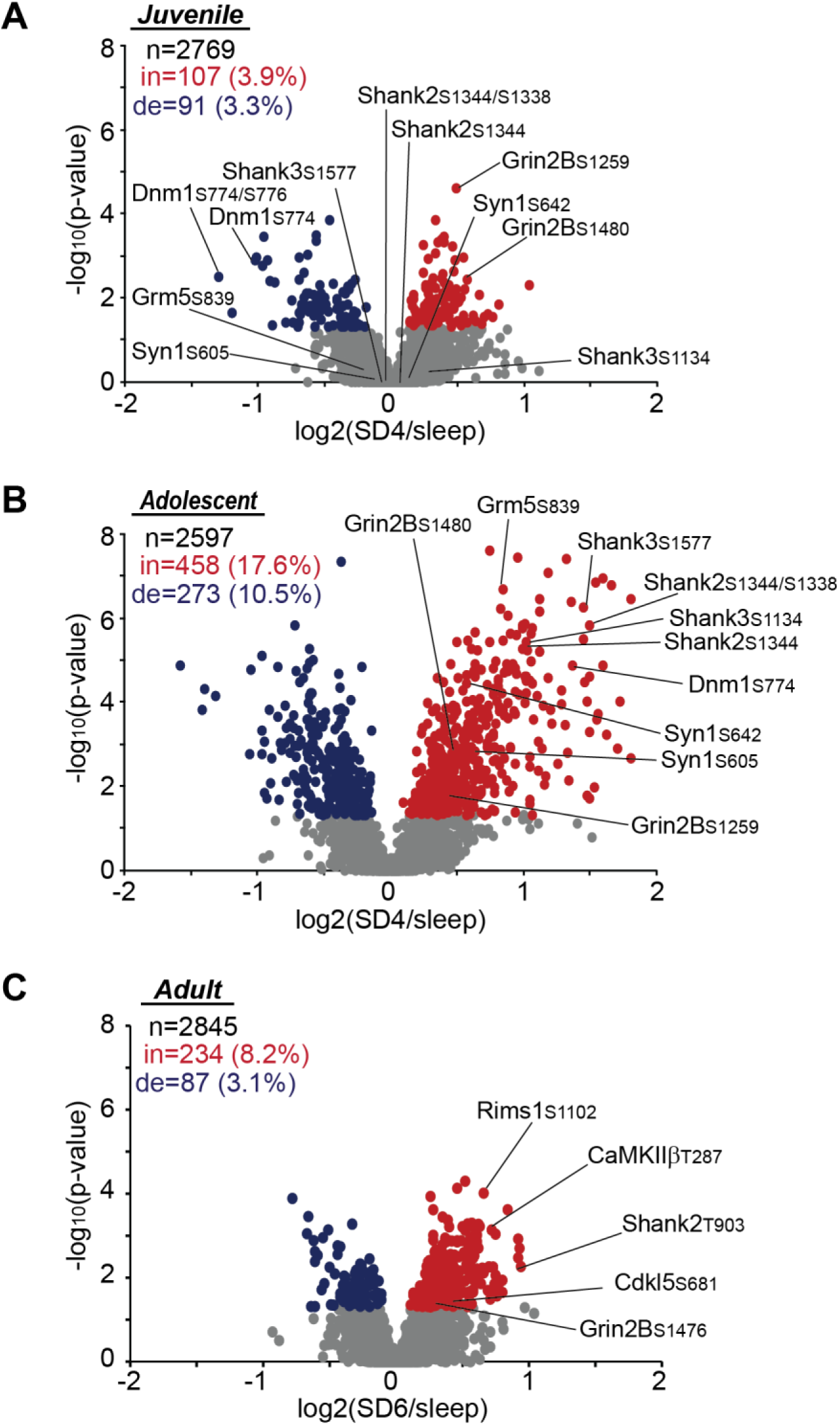
Analysis of previously identified sleep need index phosphoproteins (SNIPPs). (**A-C**) Volcano plots depicting phosphopeptides quantified from previously identified sleep need index phosphoproteins (SNIPPs) from juveniles (A), adolescents (B), and adults (C). SNIPPs display an asymmetric increase in adolescents and adults as previously described, whereas juveniles display a largely symmetric response with similar numbers of SNIPP phosphorylations increased and decreased.

## Notes

### Competing Interest Statement

The authors have declared no competing interest.

### Summary of Updates

Updated version includes new introduction and discussion. New data included in figure 1. Revised presentation and interpretation of results presented in Figure 1 and 4. New supplementary material presented in new Fig. S4.

